# Crop improvement can accelerate agriculture adaptation to societal demands and climate change

**DOI:** 10.1101/2023.09.19.558447

**Authors:** Carlos D Messina, Lucas Borras, Tom Tang, Mark Cooper

## Abstract

A key question today is how to harmonize future crop improvement efforts for regenerative cropping systems that can mitigate further environmental degeneration and improve societal adaptation to climate change. Here we show that the US corn-belt based maize improvement system has been adapting to the changing climate. Analyses of the longest running field experiment (1990-2021) designed to quantify yield gains (37 hybrids sold from 1930 to 1990) demonstrate that rates of genetic gain were always positive and have increased over time (from 70 to 150 kg ha^-1^ y^-1^ in 1990 and 2021, respectively). Between 1930 and 2021 for the May through October period total rainfall for the U.S. corn belt increased 113 mm, and daily temperature amplitude decreased 1.3°C. At the same time, farmers modified their farming practices, helping modern hybrids to out-perform their older counterparts by a larger degree. In contrast to the conclusions reached by other observational studies, genetic gain estimates demonstrate that maize breeding and the production system are adapting to modern scenarios. Climate change is commonly linked to fears of food insecurity, but new cropping systems capable of providing food while regenerating resources such as water, the circularization of nutrients, and reduction of greenhouse gas emissions are possible.

## Introduction

It is imperative to take actions to prevent net negative and inequitable impacts from climate change on agriculture, human health, the environment, and society. Various studies (*1-4*) sought to answer connected questions about the plausible impacts of climate change, genetics, and agronomy on crop yield productivity and food production; answers to these questions can inform research and policy decisions. A call for the development of traits, genotypes, and cropping systems for future climates (*5*) implies the need for implementing new dedicated gene discovery and breeding efforts. Calls for action focused solely on observational evidence of a non-ergodic system to improving stress tolerance in crops to biotic and abiotic factors are likely premature. Rather, we should seek to answer more fundamental questions: can current breeding systems continuously create genotypes adapted to a changing and uncertain climate? and, if so, how to harmonize future crop improvement efforts for regenerative cropping systems that can mitigate further environmental degeneration (*6*)? With a mounting pressure from society to act, answering these questions are of outmost importance to focus resources to adapt to the likely devastating impacts of climate change in current and future food security and social unrest if we make the wrong decisions.

Observation studies of on-farm crop production data are commonly used in attempts to deconvolute time series for yields that are the outcome of dynamical systems where genotype, environment, and agronomy (e.g., irrigation, agrochemicals, fertilizers) interact (Fig. 1) (*3, 4, 7, 8*). This approach entails risks of attributing causality to the incorrect determinants of yield improvement. In the absence of metadata, misinterpretation of causality can misguide societal investments in interventions and corrective technologies to combat the impacts of climate change for agriculture. Mechanistic crop modeling (*9, 10*) was advocated and used as an alternative approach for impact assessments (*11-14*). Studies even sought to identify ideotypes able to cope with the changing climate (*15, 16*). But mechanistic crop models are often trained for few genotypes, and assessments are conducted on how climate would affect food production in the absence of potential adaptation due to crop improvement (*12, 17, 18*).

**Figure 1.**
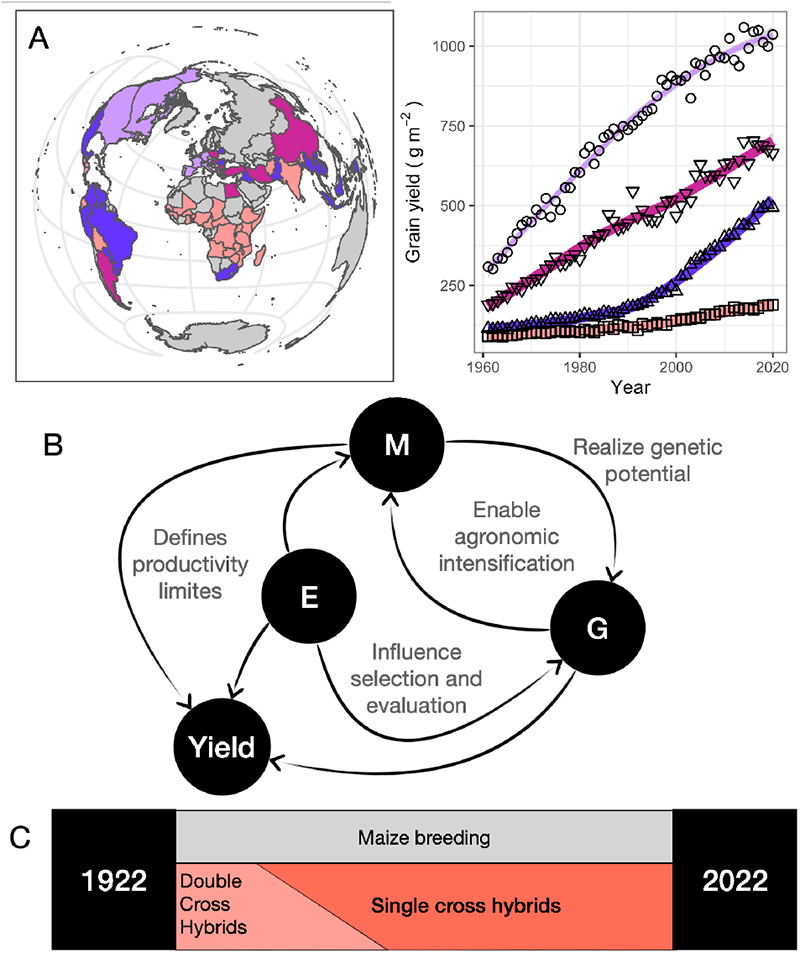
Long-term yield gain in maize worldwide (Fig 1A, colors describe regions with different farmers yield gain, FAO), results from a dynamical system where environment (E), management (M) and genetics (G) interact (Fig 1B), particularly after the 1960’s with the introduction of single cross hybrids in the U.S. corn belt (Fig 1C).

Controlled environment experiments focused on how emerging environments under climate change scenarios, higher temperature and CO_2_, would affect the physiology of crops and thus yield for a given management and genotype also used a limited set of genotypes (*19*). The integration of crop model assessments with empirical data (*18*) confronts the same challenge. While impact assessment and adaptation studies assumed that genotypes would be available as part of the toolset for climate change adaptation (*20*), it was only recently that scientists began to analyze the problem using a quantitative genetics framework (*7, 21, 22*). To date there have been no reports of long-term studies that experimentally considered genotype, management, and environments over a long period to sample the effects of climate change.

Such studies provide the experimental foundation to estimate contributions of genotype, management, environment and their interactions, and uncertainties in the determination of crop adaptation to a changing climate. Here we report results from such a long-term study for maize in the US corn-belt.

### Maize breeding in a changing climate

Concurrently with maize breeding for the U.S. corn belt (Fig 1C), the climate experienced drastic and steady changes in rainfall, minimum temperature, and amplitude temperature (Fig 2A) (*23*). In the U.S. corn belt, the rainfall for the May through October period increased by 113 mm and daily amplitude temperatures decreased by 1.3°C between 1930 and 2021. These variables impact directly and indirectly maize yields and how maize crops and genotypes interact with agronomic management (*1, 24-27*). Agronomic intensification was made possible by genetic improvement of crops to withstand plant competition and allow higher stand densities, which in turn helped harvest more water, radiation, and nitrogen (*28-31*). Farmers quickly adopted genotype (Fig 2B) and agronomic technologies (amount and type of inputs) to realize the environmental potential of their fields. Today, farmers use higher fertilizer rates and apply advanced pesticides at planting and during the crop cycle (Fig 2C-E). Field tile draining and machinery have considerably changed the capacity to reduce flooding risks and have more uniform crop stands under reduced tillage systems. Genotypes released by seed companies are continually tested on-farm in this dynamic cropping system (Fig 2B) (*33*).

**Figure 2.**
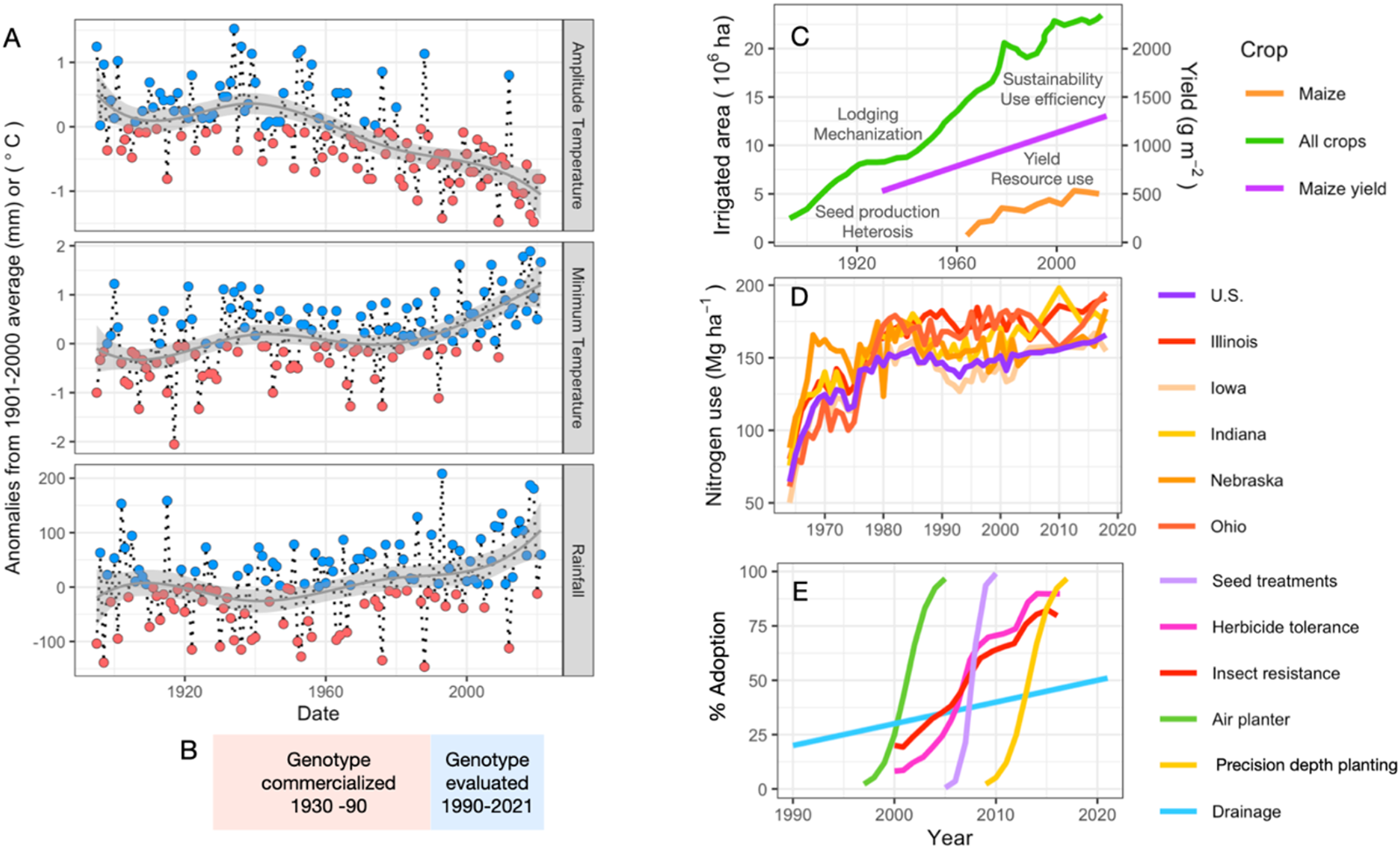
Evolution and climate variability (Fig 2A) during periods of genotype development and evaluation in field experiments (Fig 2B) designed to estimate rates of genetic gain, and concurrent changes in farming technologies such as irrigation (Fig 2C), nitrogen use (Fig 2D), seed treatments and transgenics for protecting maize crops from herbicide and insect damage (Fig 2E).

The Pioneer-Corteva maize breeding program is the longest running continuous selection experiment at commercial scale in monocots in the world. The outcomes of this program provide a unique opportunity to inform society how to approach breeding crops for future climate change. Over 100 years of maize breeding, yields increased at an average rate of 8.6 g m^-2^ y^-1^ (Fig 1C) (*2*) despite changing breeding objectives that ranged from selecting standing plants to harness mechanization technology, to select for yield to capture the benefits of fertilizers and irrigation (*1*) (Fig. 2C-E). From the 1970’s, yields for mechanized agriculture became the focus, triggered by the abundant availability of nitrogen fertilizers and irrigation in the western U.S. corn belt. Throughout this 100-year breeding journey yield potential increased continuously to produce more yield per available water, nitrogen, and solar radiation (*28-32*).

### Intervening breeding systems

What interventions are needed to maintain genetic gain in current and future climates? Using a 30 year-long continuous maize experiment conducted in the Central US corn-belt we show the capacity of a commercial maize breeding system to continuously generate new hybrid genotypes adapted to a changing climate and farm management (Fig 2). A unique feature of the data set is that a sequence of genotypes developed from the 1930s to 1990s was consistently evaluated over the 32-year period from 1990 to 2021 (Fig 3A). Therefore, it was possible to evaluate how changes in the agricultural environment from 1990 to 2021, a period where effects of climate change have been documented (Fig 2A), impacted the genetic gain (grain yield change per year attributed to genetics, Fig 3B) that was achieved by the breeding program over the preceding period from 1930 to 1990, a period when the climate also changed (Fig 2A). The 37 hybrid genotypes commercialized between 1930 and 1990 were all grown in each of the 32 years of experimentation, and within each year they were grown at multiple locations at three plant densities to represent a range of on-farm management and environments each year (*1*) (Fig 3A, Fig S1). Because rates of genetic gain could be estimated by year and agronomic management (Fig 3B) it was feasible to quantify how these rates varied over the 32-year period (Fig 3C). Counter to expectation (*3, 4*), rates of genetic gain were always positive (Fig 3C). The most significant finding of this study, however, was that the rate of genetic gain for the hybrid genotypes, developed during the period prior to the commencement of the experiment, consistently increased over the 32-year period. The genotype yields were consistently higher under modern climate and farming technologies (Fig 3C), and this was more evident under the highest plant density; genetic gain is accelerating and not decreasing. Some of the attributes that may confer improved adaptation to the changing climate include more uniform canopies, more growth, and a higher proportion of plant growth allocated to the ear, the reproductive structure bearing the harvested kernels contributing to commercial crop yield (*30, 32, 34*). This long-term experimental result demonstrates that commercial breeding can not only produce genotypes that keep pace with the changing climate and farming technologies but also show synergistic results with them. Provided breeding objectives remain unchanged it is apparent no interventions are needed.

**Figure 3.**
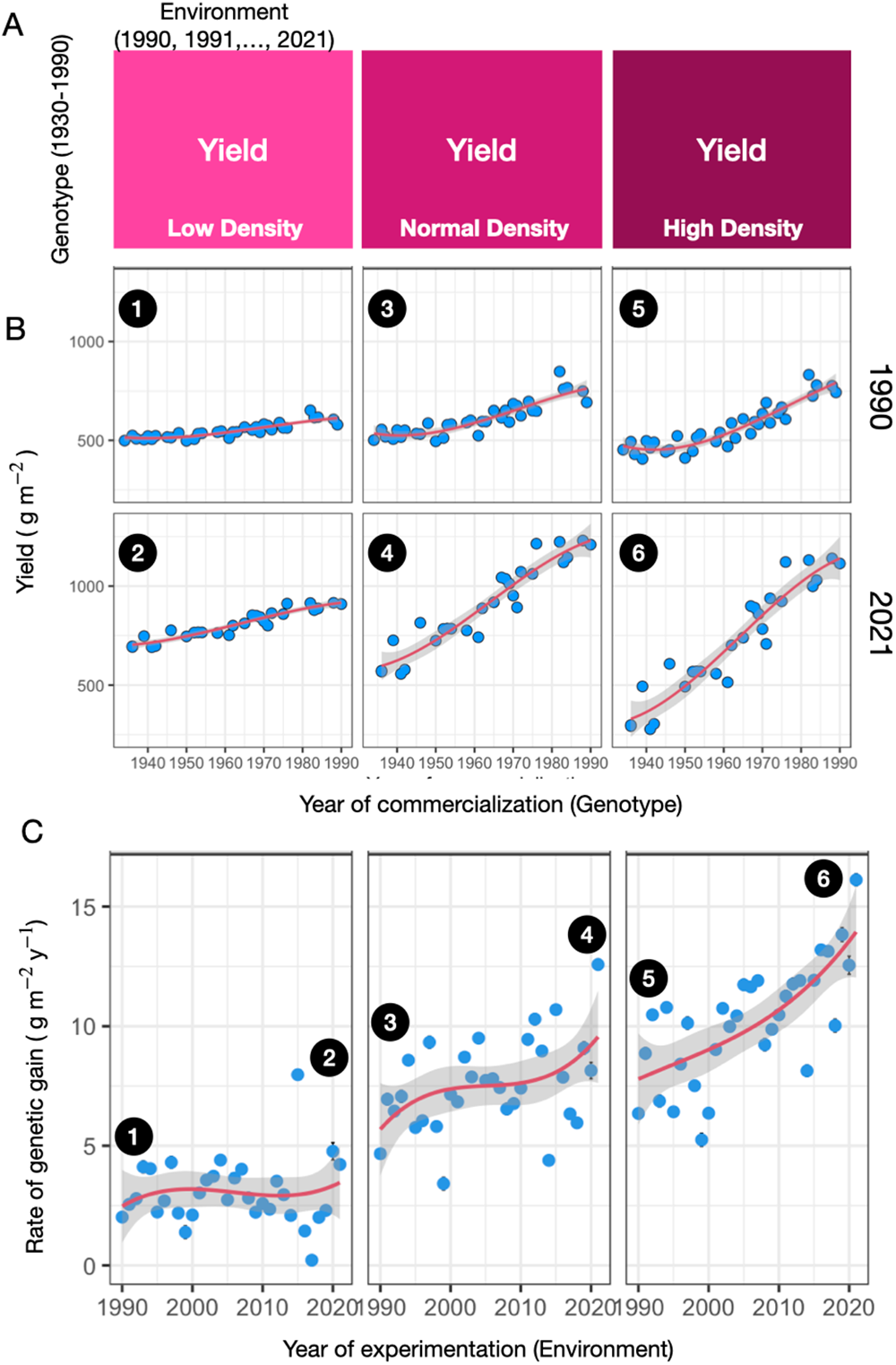
The experimental design used to evaluate genetic gain (Fig 3A) included year (n=32), planting density (n=3), and genotype (n=37), which enables without confounding estimating the genetic gain as the slope from the regression between yield of genotype against the year of commercialization (Fig 3B), and the rate of change in genetic gain from the regression of genetic gain with respect to the year of experimentation and plant density management (Fig 3C). Genetic gains are always positive and increase with increasing planting density and year of experimentation. Numbers in Figs 3A and 3B indicate the correspondence of genetic gain by year and densities.

### Breeding for modern societal needs

While new maize genotypes are more adapted to modern cropping systems (Fig 3), their use increased the cost of externalities (*35-37*). It is therefore prudent to re-consider breeding objectives to create genotypes that can contribute to modern societal needs. Breeding maize for climate change is about the capacity to maintain genetic gain for yield (Fig 3C) but doing so while reducing GHG emissions, regenerating soils and aquifers, and circularize nutrients use (*6, 38, 39*). Modern predictive breeding technologies that fuse mechanistic understanding of plant biology and quantitative genetics (*40*) provide the framework based on abductive inference to test potential genotype x management x environmental synergisms to help maintain genetic gain. Selecting the right combinations can create opportunities to regenerate aquifers by water use reductions, for example. More yield per unit of water is attained by using modern maize (Fig 4A), where the overall water use is reduced due to the lower land demand to maintain maize grain production. Similar approaches can be exemplified with other farming inputs, modern genotypes use nitrogen fertilizers more effectively for grain production and exploitable gaps for higher sustainable resource use are still available as field experimental results from modern cropping systems show (Fig 4B).

**Figure 4.**
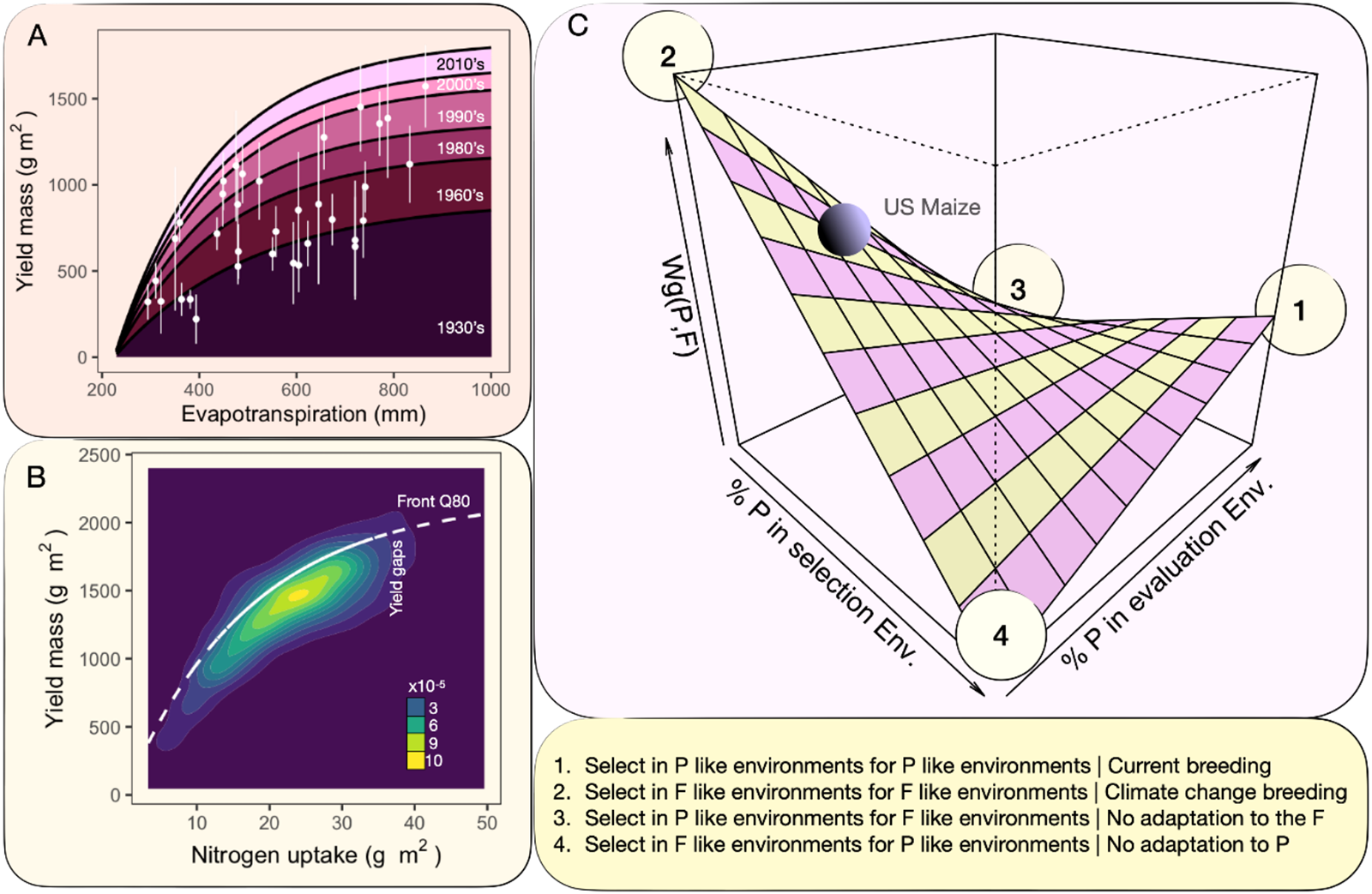
Options to combat and adapt to climate change by developing and applying genotype x management technologies: (Fig 4A) water productivity increase with genetic progress for yield represented by various groups of genotypes including open pollinated varieties commercialized in the 1930’s until single cross hybrids carrying transgenes for crop protection against insects and herbicides commercialized in in 2010’s (means and square root of the genotypic variance shown for 35 experiments) (*6*), (Fig 4B) yield gap analysis model to improve nitrogen productivity that can include choice of genotype and management variable environments (density map; n=2670 hybrid x fertilizer dose experiments), and (Fig 4C) quantitative genetics framework to inform breeding strategy for climate change adaptation based on the covariance between selection and evaluation environments (present: P and future: F) as a function of frequency of environment types.

### Improving aim at climate change

Crop breeding has been a highly impactful technology for maize yield gain worldwide (Fig 1A) (*1*). By understanding trait contributions to improved yield performance for the environmental changes that are associated with climate change we can aim at breeding for future environments under climate change by harmonizing current and future breeding efforts to maintain rates of yield gain in the present and the future on-farm environments for maize and other crops (*6, 22*). The correlated response form of the breeder’s equation (*41, 42*) provides a framework to predict genetic gain outcomes of breeding programs in future environments under the influences of climate change based on the present testing regime and breeding goals,

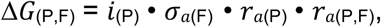

where 𝑖_(P)_ is the selection differential informed by the analyses of data collected in today’s (P = Present) trials, σ_*a*(F)_ is the additive genetic variation for the traits within the F = future target population of environments, 𝑟_𝑎(P)_ is the prediction accuracy for the trait(s) of interest estimated using the marker, phenotype and environmental information content in the training experimental sets available at present, and 𝑟_𝑎(P,F)_ is the genetic correlation between the additive genetic effects estimated for traits measured in the present and future environments. This genetic correlation between present and future environmental conditions results from the type of genotype x environment interaction and the frequency of sampling of environments that resemble current and future environment types (Fig. 4C). In the presence of cross over interaction, oversampling environments that resemble a climate change scenario may be conducive to a reduction in genetic gain and increased food insecurity today; the opposite holds if we don ‘t take efforts to consider possible changes in F (*43*). Models that link biophysics with quantitative genetics (*39, 40*) and simulation as in prior studies that used single genotypes and current and future climates (*11*) can be used to estimate 𝑟_𝑎(P,F)_. These studies must be augmented by empirical sets created using long-term field trials to monitor genetic progress and evaluate whether the structure of the breeding trials conducted in the present are predictive of future performance in a way that can keep pace with the changing performance landscapes (*44, 45*).

The structure of genotype x environment interactions and the frequency of occurrence of environment types may be conducive to compromise nutritional security in the present and/or the future (Fig 4C). Cycles of crop improvement can vary from 2 to +20 years in food crops, and depending on this time span, society will have 15 to less than 3 opportunities to create improved cultivars adapted to 2050 target population of environments. There is urgency to design and implement sound crop improvement strategies. The results presented here strongly suggest that the breeding community needs to engage in the dialogue about crop adaptation in response to climate change in multiple decision and policy forums. Currently the connection between the crop breeding community and the intergovernmental panel is dysfunctional or non-existent.

Here we report results for a single crop, which begs the question, where do we stand with other major staple food crops in other parts of the world?

## ACKNOWLEDGEMENTS

MC is supported by the Australian Research Council Centre of Excellence for Plant Success in Nature and Agriculture (CE200100015). Authors thank J. DeBruin for sharing data to build Fig. 4B.

## Supplemental Material

### Materials and Methods

A total of 103 experiments were conducted from 1990 to 2020 in the US mid-west (Fig. S1). All experiments had 37 hybrids sown at three stand densities (3.4, 5.9, and 8.4 plants m^-2^, represented as low, normal, and high stand densities, respectively) in two row plots of 5.5 meters long. Entire plots was harvested using research commercial combines, and grain weight was adjusted to report yield at 14.5% grain moisture. Fig. S1 describes the distribution of experiments across the region, and Table S1 presents the list of hybrids tested with their market release year. Hybrids were all selected based on their historical relevance, being highly sold commercial products and well known by farmers in the region over time (covering from 1930 till 1990, when the experiment started). Experiments showed no trend in sowing date over the 30 years of testing, and represented the timing that is commonly used by farmers in the region.

Weather was extracted from NOAA (https://www.ncei.noaa.gov/access/monitoring/climate-at-a-glance/regional/time-series/361/pcp/6/10/1895-2022?base_prd=true&begbaseyear=1901&endbaseyear=2000&filter=true&filterType=loess, checked 27-3-2023), and anomalies calculated for the reference period 1901-2000. Fertilizer data usage in the US was extracted from USDA (https://www.ers.usda.gov/data-products/fertilizer-use-and-price/documentation-and-data-sources/, checked 27-3-2023). U.S. average is an average of state application rates weighted by state planted acres receiving fertilizer. Tile drainage increase by 1% year^-1^ described in Fig. 1C was extracted from https://farmdocdaily.illinois.edu/2019/08/use-of-tile-2017-us-census-of-agriculture.html (checked 27-3-2023).

**Fig. S1:**
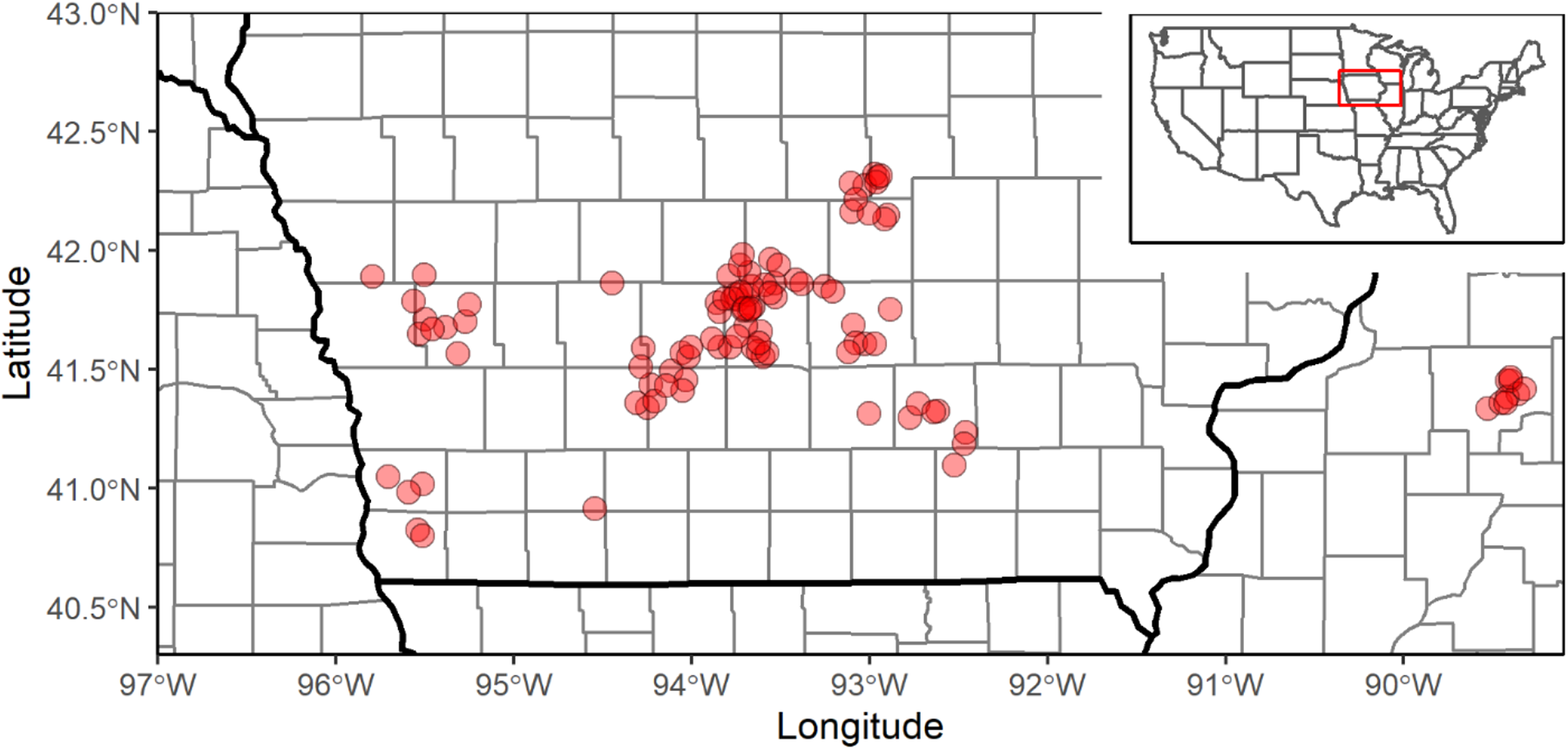
Distribution map of the 103 trials conducted from 1990 till 2020 across the US midwest. Inset shows the highlighted area in a US map.

**Table S1:**
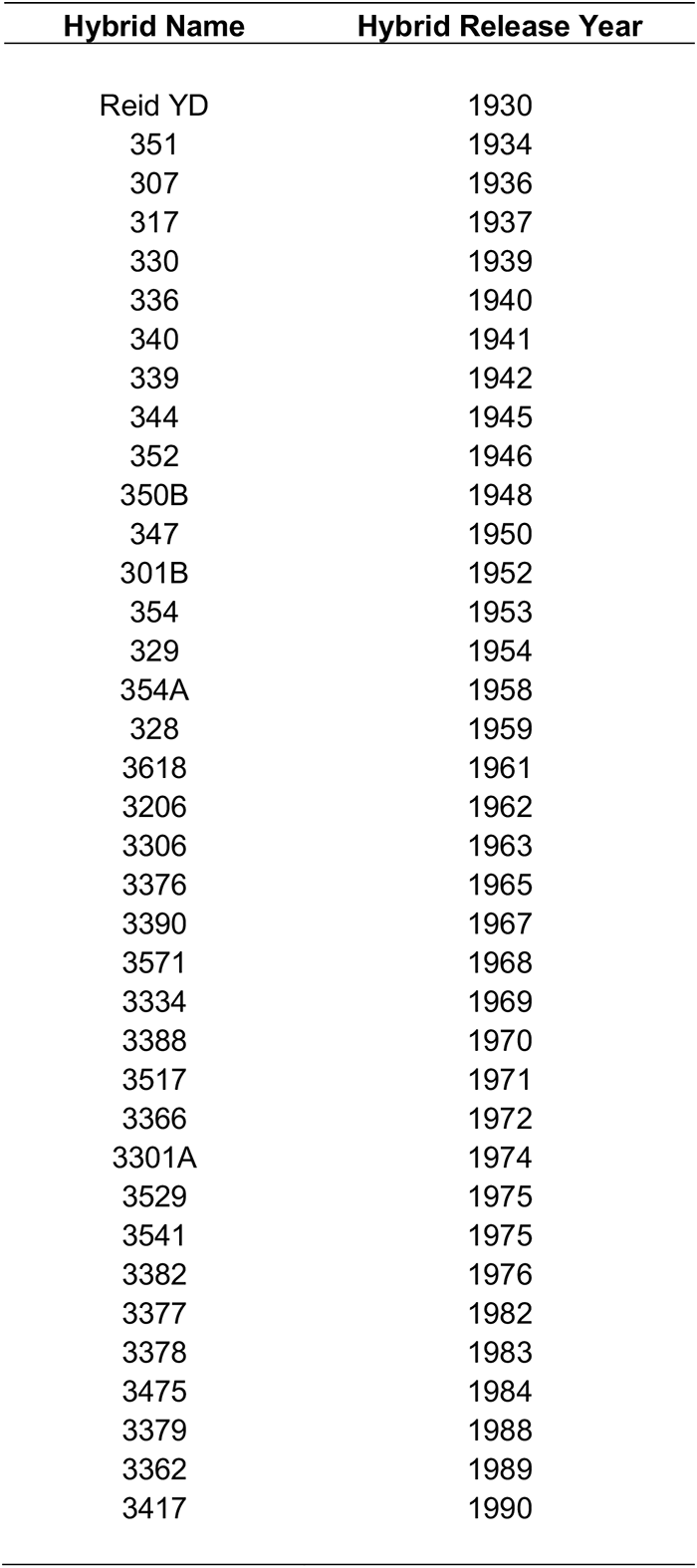
List of commercial hybrids tested from 1990 to 2020, together with their release year.

